# Style transfer with variational autoencoders is a promising approach to RNA-Seq data harmonization and analysis

**DOI:** 10.1101/791962

**Authors:** N. Russkikh, D. Antonets, D. Shtokalo, A. Makarov, Y. Vyatkin, A. Zakharov, E. Terentyev

## Abstract

**Motivation:** The transcriptomic data is being frequently used in the research of biomarker genes of different diseases and biological states. The most common tasks there are data harmonization and treatment outcome prediction. Both of them can be addressed via the style transfer approach. Either technical factors or any biological details about the samples which we would like to control (gender, biological state, treatment etc.) can be used as style components.

**Results:** The proposed style transfer solution is based on Conditional Variational Autoencoders, Y-Autoencoders and adversarial feature decomposition. In order to quantitatively measure the quality of the style transfer, neural network classifiers which predict the style and semantics after training on real expression were used. Comparison with several existing style-transfer based approaches shows that proposed model has the highest style prediction accuracy on all considered datasets while having comparable or the best semantics prediction accuracy.

**Availability:** https://github.com/NRshka/stvae-source

**Supplementary information:** FigShare.com (https://dx.doi.org/10.6084/m9.figshare.9925115)

## 1 Introduction

The new era of modern life sciences has begun with the development of high throughput nucleic acid sequencing methods – new generation sequencing (NGS) techniques. The amount of current genomic and transcriptomic data is tremendous and grows exponentially. The single cell sequencing methods enabled even more detailed description of a transcriptomic landscape that allowed to decipher the very complex nature of cellular subtypes, to analyze their developmental patterns and ancestry (Saliba *et al.*, 2014; Stark *et al.*, 2019).

However, current NGS data is highly fragmented due to different sources of technical variation associated with particular NGS platforms, sample acquisition and preparation procedures, subsequent analysis steps etc. The costs of transcriptomic experiments are still high and thus the really big datasets comprising thousands of samples are still rare. One of the most frequent tasks in transcriptomic data analysis is the identification of potential biomarker genes for various diseases and conditions. In most cases the researchers operate with data comprising from tens to hundreds, and, in rare cases, thousands of samples and the tens of thousands of genes or individual transcripts. The extremely high dimensionality and complex mutual interdependencies make it difficult to achieve reproducibility in prospective studies. The problem is the insufficient volume of any single training dataset, an excessively large number of influencing factors and the lack of knowledge about the structure of molecular genetic systems. Thus, there is an urgent need in methodological approaches capable to analyze heterogeneous and limited datasets of high dimensionality, suffering from technical noise and different kinds of batch effects. One of the available options is to harmonize the quality control procedures and the data analysis pipeline to make the resulting gene (transcripts) expression values more comparable. One of the best examples of this approach is DEE2 – Digital Expression Explorer 2 (DEE2) (Ziemann *et al.*, 2019) – a repository of uniformly processed RNA-seq data obtained from NCBI Short Read Archive. There are also other examples: ARCHS4 the massive collection of uniformly processed murine and human public transcriptomic datasets (Lachmann *et al.*, 2018), recount2 (Collado-Torres *et al.*, 2017) etc. However, the most important task in transcriptomic data harmonization is the correction of batch effects and in general it remains unresolved.

Currently it is widely accepted that gene expression profiles of the living cells resulted from a complex mixture of different biological processes and technical parameters. At the moment, there were several attempts to model this kind of data as combinations of certain low-dimensional representations corresponding to various biological pathways and conditions (Xu *et al.*, 2019). In this work we test the hypothesis whether these attributes could be reasonably and controllably changed in silico using the deep learning models.

We believe that both batch effect correction and treatment outcome prediction tasks may be addressed with a style transfer approach. In the first case, we consider each batch as a style and transfer all of the samples to the same style to eliminate batch discrepancy. In the latter case, we consider “Treatment’ and “Non-Treatment” conditions as styles and transfer the style of the desired samples from “Non-Treatment” to “Treatment” and vice versa. Please note that we use the term “batch” here and further in the text in purely biological sense, outside of the stochastic gradient descent algorithm scope. For the latter, we use the word “minibatch”. In this work we applied adversarial decomposition methodology to disentangle the sources of variation in single cell RNA-Seq data in order to be able to transfer its style. In our approach we used no prior dimensionality reduction as it makes strong assumptions about the data. For example, PCA tries to maximize the variance in projected dimensions and in case of heavy outliers and non-symmetric distribution the result becomes unstable at least if one doesn’t apply the robust covariance estimates. Another common problem is that top PCs often extract the technical variation. Besides, we assume it’s unlikely that biological states can be modelled by simple linear combinations of some low dimensional basis vectors since different sorts of non-linear relations are common for gene regulation circuits e.g., logical XOR patterns, various feedback loops and conditional dependencies etc. Given the highly hierarchical modular organization of cellular regulatory pathways and the clonal nature of the cells, deep neural network-based approaches seem to be the most feasible for the tasks involving gene expression.

## 2 Background

In recent years, there have been plenty of studies applying deep learning to gene expression data analysis. Speaking of architecture, they can be roughly subdivided on GAN-based, autoencoder-based and Bayesian deep learning-based. With respect to the tasks, those models may serve for denoising, missed values imputation, dimensionality reduction (and further analysis of obtained low-dimensional embeddings), data harmonization and in silico treatment outcome prediction. For example, an approach with deep generative modeling for scRNA-Seq data normalization and domain adaptation was recently proposed in (Xu *et al.*, 2019; Johansen and Quon, 2019). Another approach to gene expression data modelling with autoencoders was presented in (Gold *et al.*, 2018) – the authors induced the sparsity of network weights by connecting only the genes from the same functional group to the same hidden neuron. This is a step towards interpretability of autoencoder models. Authors of (Way and Greene, 2018) and (Grønbech *et al.*, 2018) successfully used variational autoencoders (VAE) as a non-linear dimensionality reduction method for gene expression data from different cancer subtypes and cell types, respectively. In (Eraslan *et al.*, 2019) the autoencoder with zero-inflated negative binomial (ZINB) likelihood loss was effectively used as a denoising tool on gene expression data. In (Amodio, M. *et al.*, 2019) the sparse autoencoder-based approach was proposed for clustering, imputation and embedding of the single cell transcriptomic data. In (Wang, Dongfang & Gu, Jin. 2018) the VAE-based model with additional layer accounted for zero-inflation was proposed for the single cell data imputation. The approaches with style transfer involved are usually aimed to address the imputation and embedding tasks. Originally this type of transformation was mostly applied to adopt the style of fine art paintings to generic images (Gatys *et al.*, 2015). A large group of image style transfer frameworks uses the pretrained models to extract image descriptors in order to build the transfer objective. Due to the absence of such pretrained models for gene expression data, we made use of adversarial approach: separating the features into style and semantic groups using the discriminator network. Learning the representations independent of domain with the help of discriminator using gradient reversal layer was proposed in (Gatys *et al.*, 2015). The adversarial decomposition strategy was successfully applied to style transfer of texts, for example in this work (Romanov et al., 2018). In our work, we also used cycle-consistency loss for style transfer, which was proposed in (Zhu *et al.*, 2017). The same technique used in domain adaptation for image segmentation tasks can be found in (Hoffman *et al.*, 2017).

In gene expression analysis the style transfer methods are still not widespread and certain frameworks and models were developed only recently and applied mostly for single cell transcriptomic data. In some of them, transfer is done by latent vector arithmetics, i.e. a vector connecting the averaged latents of initial and target cell groups is added to the initial latents and then the shifted latents are decoded back to the original gene expression space. Such an approach with GAN-based latent space mapping was proposed in (Ghahramani *et al.*, 2018) and with VAE-based model in (Lotfollahi *et al.*, 2019a). Yet another GAN-based approach (Targonski *et al.*, 2019) uses the adversarial attack technique to perturb gene expression profiles to fool the pretrained category classifier, which also may be considered as a style transfer. Another autoencoder-based approach with shared decoder and one decoder per style was implemented in recent work by Johansen *et al.* (Johansen *et al.*, 2019). Among the solutions based on Conditional Variational Autoencoder one should also mention the recent work by Lotfollahi *et al.* (Lotfollahi *et al.*, 2019b) where the authors used maximum mean discrepancy (MMD) loss to achieve better disentanglement of features related to data style and semantics. Inspired by these results we decided to study if different components of gene expression data variance can be disentangled using Convolutional Variational Autoencoder (CVAE) based techniques and adversarial decomposition and if expression with transferred style obtained via such disentangled representation might be of interest from a biological point of view, e.g. for in silico treatment outcome prediction, data augmentation with semisynthetic samples etc.

## 3 Methods

### 3.1 Datasets

#### 3.1.1 The Murine Cell Atlas (scMCA)

This dataset comprising numerous murine single cell gene expression profiles was produced with cost-effective high throughput Microwell-seq platform (Han *et al.*, 2018), that allowed to analyze over 400,000 single cells from 51 mouse tissues and organs extracted from several animals at varying physiological conditions. The original scMCA data contains gene expression profiles for over 800 major murine cell types. The detailed annotation was provided by the authors for over 200,000 single cells. A detailed description of the data can be found in the original paper (Han *et al.*, 2018) and online. This dataset was selected due to the following major reasons: (1) it contained the huge amount of data obtained with a consistent methodology by the same research group thus presumably making the technical dispersion less profound; (2) since the samples belong to different animals, distinct organs/tissues and physiological conditions one could build a model to decompose these sources of variation.

For building the models we selected a subset of 45497 samples corresponding to single cells derived from murine mammary glands of virgin and pregnant mice and also from involution state (24395, 9737 и 11365 samples respectively). This subset was selected both due to its volume and the ease of interpretation of distinctions between conditions. The samples from lactating animals were excluded as they were clearly isolated (data not shown) and different cellular types were barely distinguishable. We kept 20% of the data (with keeping the same proportion of biological states) as a test set, and 15% of the remaining data was used for validation. The original raw gene expression counts were used as inputs. 15987 genes were taken into consideration. The gene expression tables used in our study can be found in supplementary tables (ST1). To reduce the cell type labelling complexity, we decided to switch to more general cell types categories, presuming that the expression patterns between the cells of common origin should be more consistent than those of different cellular types. The complete data annotation used in our experiments is listed in supplementary table ST2. We considered the cellular types as the element of data semantics. Among the major goals was to control the preservation of related cell types under the style transfer procedure. The original scMCA data can be found at: https://doi.org/10.6084/m9.figshare.5435866.v7.

#### 3.1.2 STARmap

The STARmap dataset was used for hyperparameters tuning and comparative testing of our model against several other approaches (see below). It contains the expression values for 166 genes in 3,700 cells from three separate biological mouse samples of the medial prefrontal cortex (Wang et al., 2018). The annotated dataset was taken from https://github.com/YosefLab/scVI-data/raw/master/mpfc-starmap.loom from the authors of scVI framework (Lopez *et al.*, 2018). Loom is a specialized file format based on HDF5 suitable for large omics datasets, containing a main data matrix and additional annotation layers. The loompy – a Python library for working with Loom data can be found at: http://loompy.org.

#### 3.1.3 Retina

The original dataset contains 27,499 cells and 13,166 genes from two batches (Shekhar *et al.*, 2016). This dataset was also used for benchmarking. We used the cluster annotation from 15 cell types and preprocessed and normalized gene expression counts provided by scVI authors (Lopez et al. 2018). The annotated dataset can be downloaded from https://github.com/YosefLab/scVI-data/raw/master/retina.loom.

#### 3.1.4 PBMC

The data was originally extracted from SRP073767 dataset by (Zheng et al. 2017). It is the scRNA-seq data from two batches of PBMCs from a healthy donor (4,000 and 8,000 PBMCs, respectively). The dataset was prepared as described in scVI paper (Lopez et al., 2018); the annotated dataset contained 12,039 cells with 3,346 genes. The dataset was used for benchmarking. The gene expression data can be downloaded from https://github.com/YosefLab/scVI-data/raw/master/gene_info.csv and the corresponding metadata - from https://github.com/YosefLab/scVI-data/raw/master/pbmc_metadata.pickle.

#### 3.1.5 IFNβ-treated PBMC

For biological validation we have also used the dataset containing control and interferon-beta stimulated PBMCs (GSE96583) (Kang *et al.*, 2018). The data was taken from scGen examples (https://github.com/theislab/scgen-reproducibility). The dataset was provided by the authors (Lotfollahi *et al.*, 2019a) as normalized and logtransformed. The data included 18,868 cells belonging to 8 cellular types and 6,998 genes in two conditions. The examples can be found at their project repository: https://nbviewer.jupyter.org/github/M0hammadL/scGen_notebooks/blob/master/notebooks/scgen_kang.ipynb.

### 3.2 Deep learning model development

#### 3.2.1 Autoencoder architecture

Our architecture is based on multiple autoencoder-related ideas. We use Conditional Variational Autoencoder (Sohn *et al.*, 2015) as a backbone for our encoder-decoder architecture: one-hot encoding of the category is fed to the decoder as well as latents. This kind of architecture makes us able to perform style transfer: after encoding of the initial expression, we can choose a target category before decoding. In order to eliminate the information about the category which represents our ‘style’ from the hidden variables, we place a discriminator over the latent representation and train it adversarially with the encoder. To further enforce the disentanglement between the style and the remaining hidden variables, we adopt the training techniques from Y-Autoencoder (Pattachiola *et al.*, 2019) which involves adding yet another head to the encoder which predicts the category and minimizing a bunch of auxiliary loss functions along with reconstruction loss. Additional hyperparameter aimed to weight a contribution of Kullback-Leibler divergence with prior distribution to the total loss as in beta-VAE (Higgins *et al.*, 2017) is added. Given the input dimension, number of layers and bottleneck size, we control the width of the i^th^ layer with the following formula:

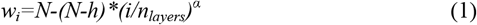

where *N* is a input dimension, *h* is a bottleneck size and *α* is a hyperparameter which controls how fast the layers width shrinks when moving from input to bottleneck.

We used nonlinearities Mish (Misra, D., 2019) and (mini)batch normalization in both encoder and decoder layers. The architecture scheme is presented on Fig. 1. Discriminator scheme is the following: *Input-FC(1024)-BatchNorm-LeakyReLU-FC(1024)-BatchNorm– LeakyReLU-FC(N_batches)*, where abbreviature *FC* stands for fully connected layers.

**Fig. 1.**
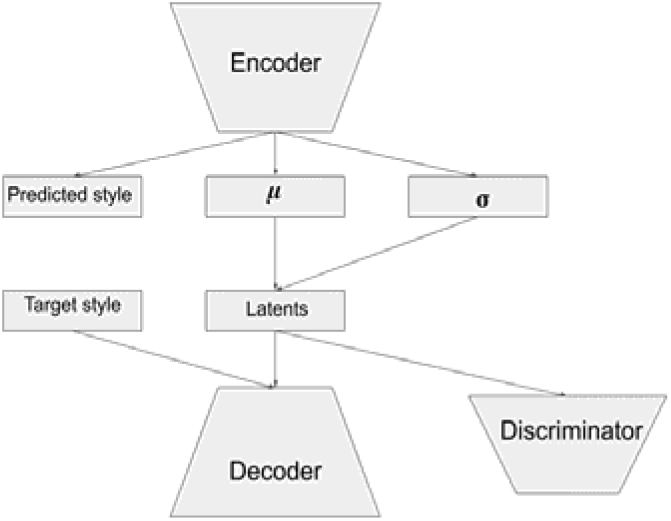
The autoencoder architecture scheme.

#### 3.2.2 Autoencoder training

For the training of our autoencoder, we used the mean squared error (MSE) as reconstruction loss function. Also, a cyclic consistency loss was used: we obtain the encodings for a minibatch, make a random style transfer, and then transfer the style back at the second forward-pass through the autoencoder.

Reconstruction loss between the values obtained this way and the initial expression is a cycle consistency loss. In order to enforce the hidden representation to contain no information about biological state, we maximized Shannon entropy of discriminator predictions as generator loss. Discriminator was trained with a log-loss objective. Auxiliary losses from Y-Autoencoders (Pattachiola *et al.*, 2019) were also minimized. Their coefficients were set equal to reduce the dimensionality of hyperparameter search.

For regularization we used the L1 weight penalty for autoencoder along with VAE-regularization. For adversarial training stabilization, we have used the gaussian instance noise (Mescheder, 2018) with variance 0.01 for discriminator. Also, gradient clipping down to unite norm was used for autoencoder and discriminator was used. To sum it up, the training of our model may be described with pseudocode shown in supplementary file SF1.

The weights for each term in the autoencoder along with number of model layers and alpha hyperparameter were tuned with random search with roughly 350 iterations on the STARmap dataset. The optimal hyperparameters were the following: cvae_beta = 2e-5; adv_weight = 0.0000001; vae_lr=0.001; num_epochs=800; n_layers = 2; scale_alpha=1.3; bottleneck_size=30; form_consistency_weight = 0.2; batch_size = 128. To get a better understanding of these hyperparameters and their roles please see the Supplement file SF1 with pseudocode.

For downstream analysis of autoencoder outputs, we substituted the predicted negative values with zero. Several experiments with ReLU activation were used as the last layer to prevent the appearance of negative outputs were conducted, but these led to poor model convergence.

#### 3.2.3 Metrics classifiers architecture and training

In order to quantitatively measure fidelity of the style transfer and semantics preservation during this procedure, auxiliary neural network classifiers were used. They are needed just to measure the performance of the frameworks on the style transfer task and did not participate in their learning whatsoever. Architecture scheme is *Input-FC(512)-BatchNorm-Mish-FC(256)-BatchNorm-Mish-FC(128)-BatchNorm-Mish-FC(OUTPUT_SIZE)*. Both classifiers were trained with Adam optimizer for 450 epochs with minibatch size 128. Learning rate was set to 0.003 for cell type classifier and 0.00001 for the style classifier. These hyperparameters were hand-picked in a set of experiments.

#### 3.2.4 Other frameworks architecture and training

1. **trVAE** (Lotfollahi *et al.*, 2019b). Implementation from https://github.com/theislab/trvaep was used. The model contained two hidden layers in both encoder and decoder with sizes of 128 and 32, respectively. Bottleneck layer size was set to 30 (as well as in all other frameworks). Hyperparameter alpha was set to 0.0001. The model was trained for 300 epochs with minibatch size of 512 with early stopping patience of 50 epochs.
2. **scGEN** (Lotfollahi *et al.*, 2019a). Implementation from https://github.com/theislab/scgen was used. Bottleneck was set to 30 neurons, all of the other hyperparameters used the default set-ting.
3. **scVI** (Lopez *et al.*, 2018). Implementation from https://github.com/YosefLab/scVI was used. All the hyperparameters used the default setting besides the number of latent variables, which was set to 30.
4. **CycleGAN** (Zhu *et al.*, 2017). Implementation from https://github.com/junyanz/pytorch-CycleGAN-and-pix2pix was used. Since this implementation was meant to be used for the im-age data, we modified the autoencoder architecture to *Input-InstanceNorm-ReLU-FC(365)-InstanceNorm-ReLU-FC(30)– In-stanceNorm-ReLU-FC(365)-InstanceNorm-ReLU-FC(OUTPUT_SIZE)* for all the datasets besides STARmap where the hidden layer had 94 neurons instead of 365 due to lower input dimensionality. The discriminator scheme was modified to *Input-InstanceNorm-ReLU-FC(365)-InstanceNorm-ReLU-FC(1)*. Dis-criminator loss function was set to binary cross entropy instead of mean squared error. All other infrastructure and hyperparameters left unchanged.

#### 3.2.5 Calibration procedure

Yet another, simple approach to validate the models is what we call a *calibration procedure*. It is designed to control that keeping the original sample style while passing the sample through the model provides less deviation of expression than an arbitrary style transfer. Namely, we take a sample, transfer its style in all possible ways and check if L2-distance between the original and decoded expression achieves the smallest value when the initial sample style is used. One may think of it as a simple rule-based classifier.

### 3.3 Biological assessment and validation

#### 3.3.1 MA-plots construction

Each point on the MA-plot is a gene. Sum of expression of each gene was calculated across all samples belonging to the particular cell type in the same state and 1.0 was added to avoid division by zero proble. The abscissa is calculated as an average of log2-transformed expression of a gene in two compared states. The ordinate is the log2 transformation of the fold change of expression between two compared states.

#### 2.3.2 Differential gene expression and gene set enrichment analysis

With scMCA data the differential gene expression analysis was performed using RPM-normalized expression counts. The statistical significance was assessed with Mann-Whitney test with multiple testing p-value correction using FDR procedure. Several cellular types were processed separately: (1) Stromal/Luminal/Alveolar cells – those functionally involved in mammary gland development and lactation and (2) Dendritic cells – antigen presenting cells that were expected to display less profound differences between virgin, pregnant and involution states. GO- and KEGG-enrichment analysis were performed with the online resource ShinyGO (v0.60) (Ge and Jung, 2018). The lists of murine genes, associated with certain GO-categories were taken from Gene Ontology Browser at Mouse Genome Informatic portal (Bult *et al.*, 2019).

With IFNβ-treated/control PBMC scRNA-Seq data the differential gene expression analysis was performed with either Mann-Whitney or Welch’s test with Bonferroni p-value adjustment. GO-terms enrichment analysis was performed with Python package goenrich (https://github.com/jdrudolph/goenrich). All the details can be found n Jupyther notebooks at our project repository.

## 4 Results

Our research was aimed to disentangle the information about the cell type and biological state in the low-dimensional representation of gene expression data. Since gene expression data is more interpretable and familiar to bioinformaticians and is also suitable for downstream analysis pipelines than low-dimensional embeddings, we paid more attention to evaluating the results of our model output expression rather than latent representation. However, we also report two metrics related to the latent representation, namely knn purity and entropy of batch mixing (Xu *et al.*, 2019). The plots illustrating

The disentanglement can be also illustrated with the following examples. Fig. 2 and Fig. 3 depict the 2D projections of the testing samples obtained with tSNE using either original gene expression values or the recovered expression obtained with our model, respectively. The samples are colored according to cell types (A) and to condition (B). One can readily see the clusters corresponding to cell types and to condition on both these plots. However, when similar visualization was built using the extracted latent representations of the samples as input (Fig. 4), there were no clusters corresponding to different physiological states, but the clusterization of cell types was still observed. We have additionally obtained the low dimensional projections of scMCA and GSE96583 with UMAP. The figures were found to be more informative and there were evident clusterizations of scMCA data points even on latents. The corresponding figures S1 and S2 can be found in Supplementary file SF2.

**Fig. 2.**
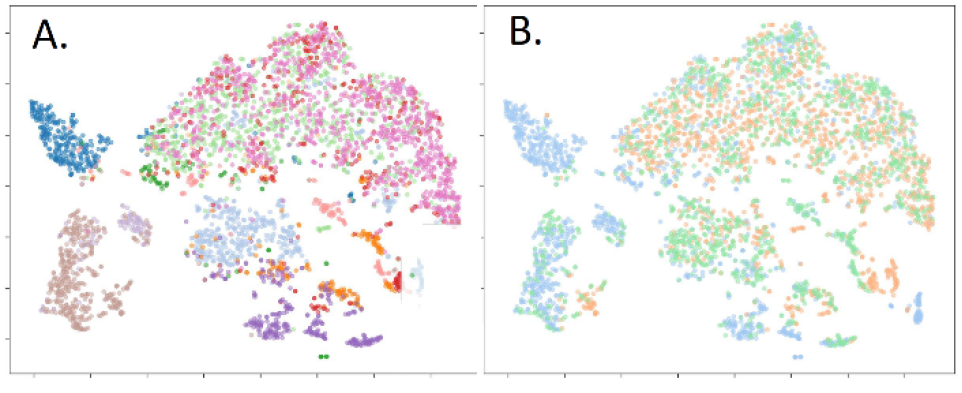
Visualization of original samples with tSNE. Raw expression values were used, samples were colored according to cell types (A) and physiological state (B). tSNE perplexity was set to 30.

**Fig. 3.**
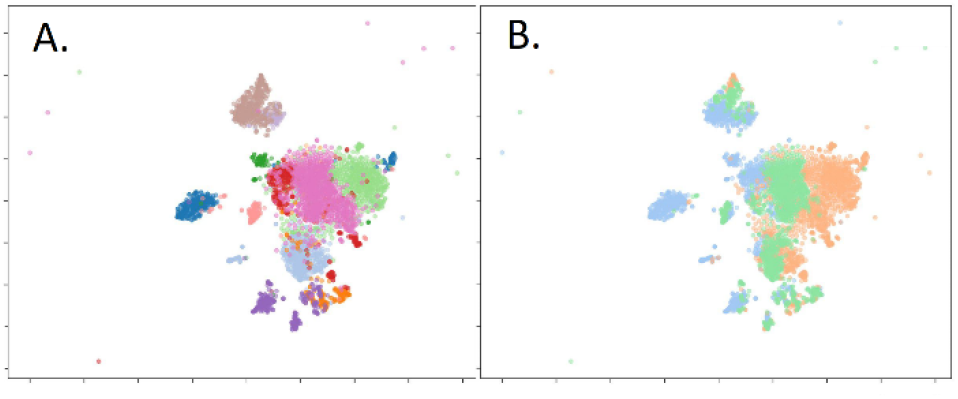
Visualization of reconstructed samples with tSNE. Gene expression values reconstructed with VAE model were used, samples were colored according to cell types (A) and physiological state (B). tSNE perplexity was set to 30.

**Fig. 4.**
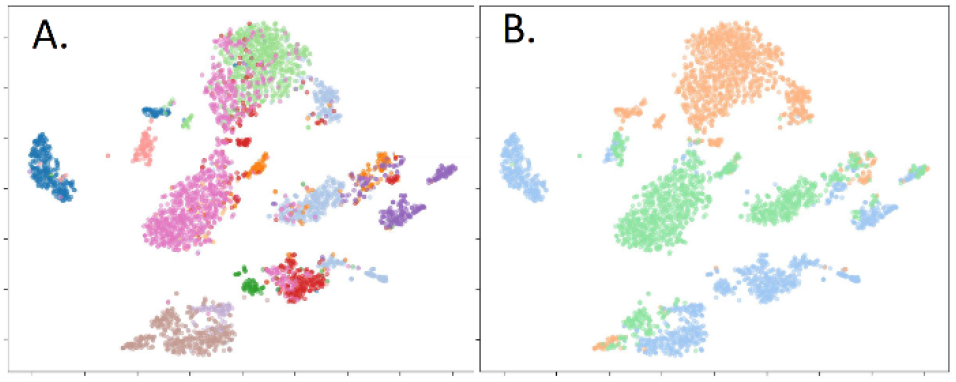
Visualization of the samples with tSNE using the learned latent representation. The latent variables of the testing samples were obtained with pre-trained encoder. The samples were colored according to cell types (A) and physiological state (B). tSNE perplexity was set to 30.

### 4.1 Style transfer validation

We expect our model to be able to switch the gene expression with n the samples in a discrete set of categories (style) while keeping all other properties preserved, e.g. changing the condition without affecting the cell type. In order to quantitatively measure that, we use neural network models which predict the style and the cell type after training on real gene expression sets. Ideally, these models should correctly predict the style the samples were transferred to and cell type predictions should remain invariant despite the style transfer. Several autoencoder-based style transfer approaches along with our model were evaluated this way with the same pretrained classifiers on several datasets with the same train/validation/test splitting. Results of the comparison are presented in Table 1.

**Table 1.**
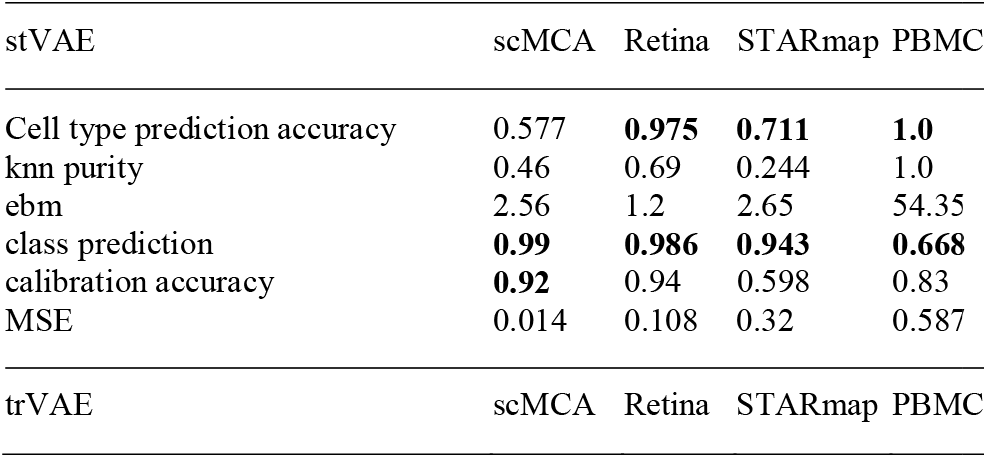
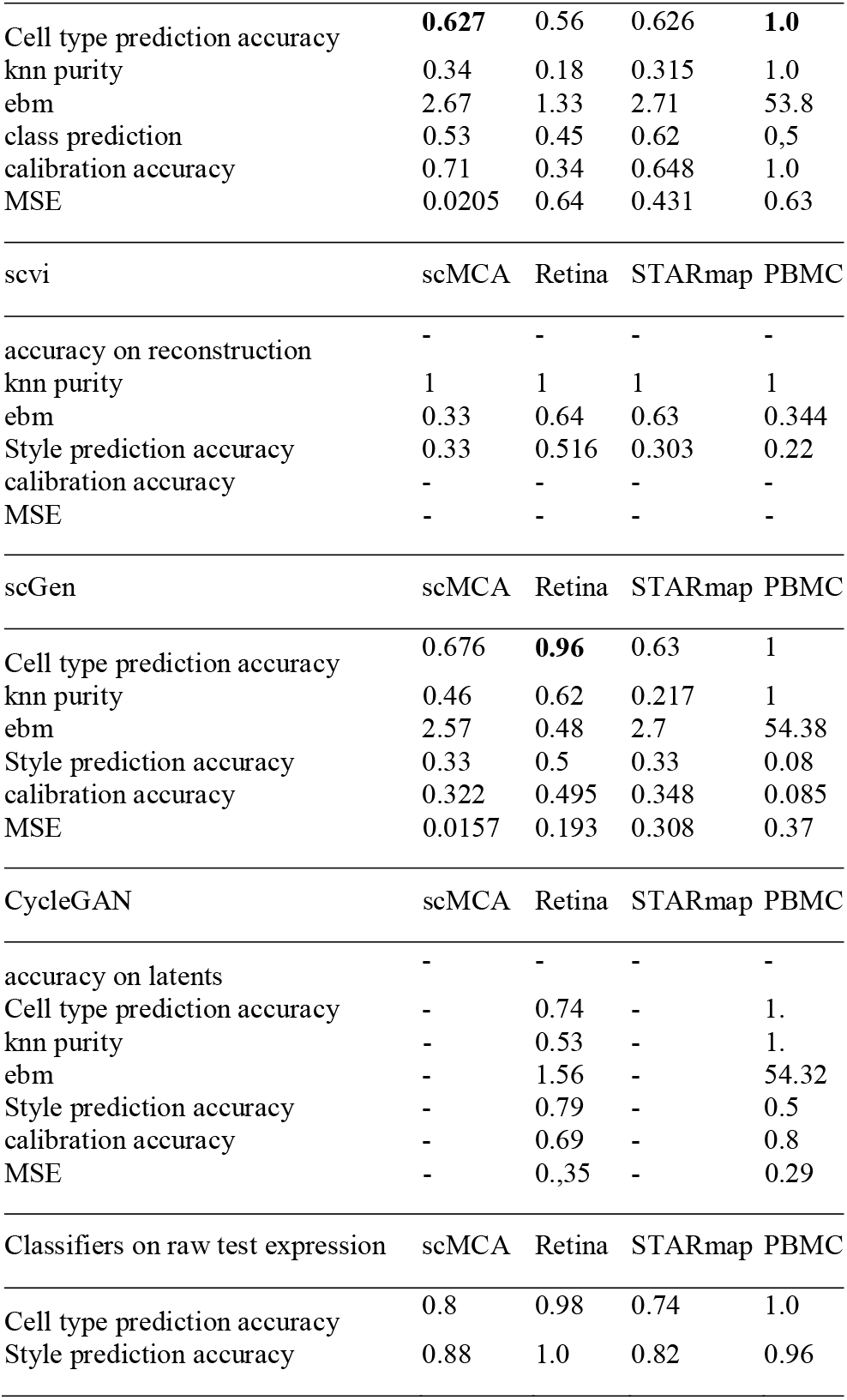
Comparison of several style transfer-based frameworks with proposed model in terms of several performance metrics using different testing datasets

It turns out that our model provides the highest style prediction accuracy on all of the datasets while providing the highest cell type prediction accuracy on two datasets and the second highest accuracy on the remaining ones. These results suggest that latent space arithmetics is not sufficient to transfer the sample style.

### 4.2 Ablation study

We have conducted the ablation study on the mouse dataset in order to analyze what parts of the model have the biggest impact on the style transfer fidelity. The parts were disabled by zeroing out the corresponding weight in the loss function. The results may be found in Table 2. The top row contains information about the loss weight value when the corresponding part is enabled. They are set to the hyperparameters found by random search on the STARmap dataset.

**Table 2.**
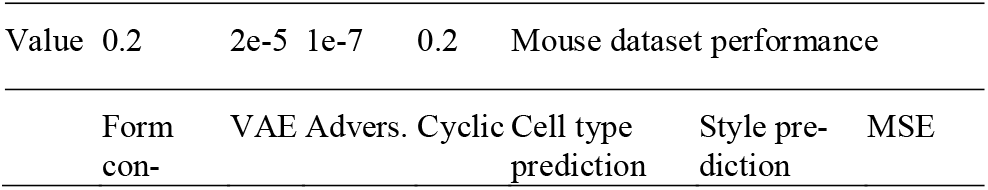
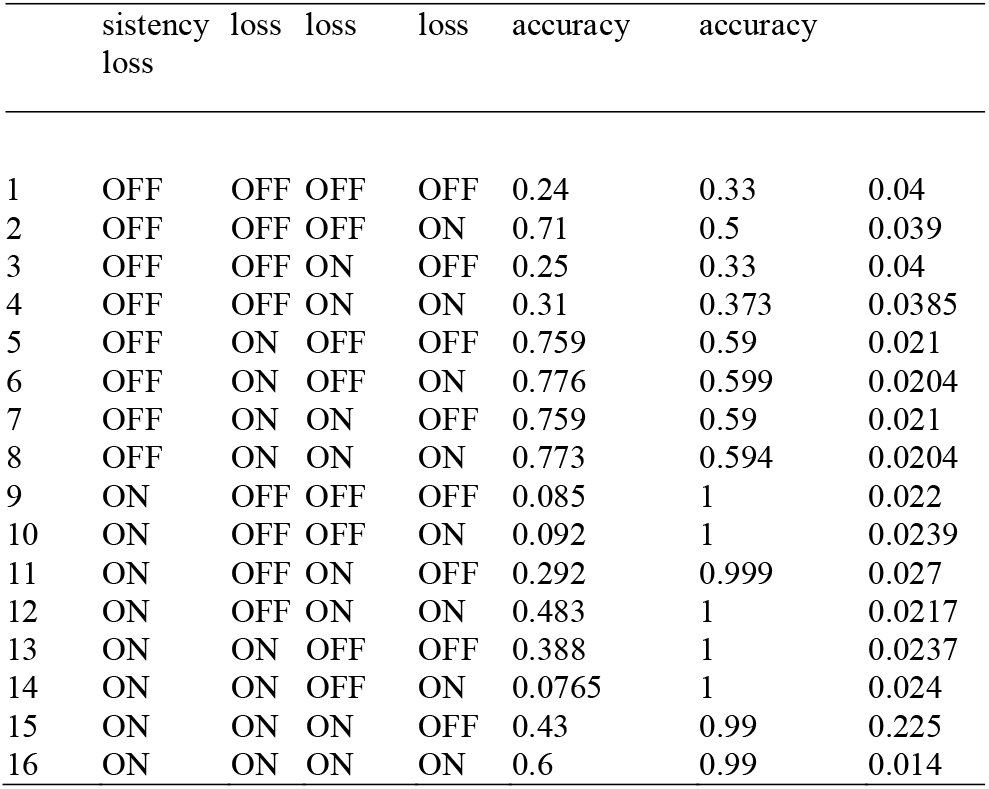
Comparison of several style transfer-based frameworks with our model with respect to different metrics on testing set

The table shows that the model benefits greatly in terms of style prediction accuracy from Y-Autoencoder auxiliary losses, sum of which we call the form consistency loss. VAE regularization is mostly responsible for cell type prediction accuracy. Both classification metrics are slightly improved by introducing the cycle consistency loss. Since the adversarial loss was given very low coefficient after the random search its impact is negligible.

### 4.3 Biological examination of gene expression changes after encoding and decoding transformation using scMCA data

The verification was performed using differential gene expression analysis and gene set enrichment analysis with GO and KEGG categories. Differential gene expression analysis was performed using RPM-normalized expression counts. The statistical significance was assessed with Mann-Whitney test with multiple testing p-value correction using FDR procedure. Several cellular types were processed separately: (1) Stromal/Luminal/Alveolar cells – those functionally involved in mammary gland development and lactation and (2) Dendritic cells – antigen presenting cells that were expected to display less profound differences between virgin, pregnant and involution states.

GO-enrichment analysis demonstrated that used data contained relevant biological signals (Table 3). When Stromal/Luminal/Alveolar cells taken from mammary glands of pregnant mice were compared against those of virgin mice, the top 100 upregulated differentially expressed genes were found to be significantly enriched with epithelium development, epithelial cell differentiation, mammary gland development GO categories. The top 200 upregulated genes were also found to be significantly associated with progesterone-mediated oocyte maturation and prolactin signaling KEGG pathways. The top 100 upregulated differentially expressed genes found with comparison of Stromal/Luminal/Alveolar cells from mice with mammary gland involution against those of pregnant animals were found to be significantly enriched with GO categories related to apoptosis, stress response and catabolism (Table 3). When similar analysis was performed using dendritic cell samples, the top 100 differentially expressed upregulated genes were found to be significantly enriched with GO categories related to defense and immune responses, cytokine production, dendritic cell differentiation etc. (data not shown).

**Table 3.** GO-enrichment analysis of top 100 differentially expressed genes observed in Stromal/Luminal/Alveolar cells in virgin vs. pregnant and involution vs. pregnant comparisons

| GO-enrichment analysis of Stromal/Luminal/Alveolar cells of top 100 genes found to be differentially expressed in samples of virgin and pregnant mice |  |  |  |  |
| --- | --- | --- | --- | --- |
| Enrichment | Genes | Total | Functional Category | GO ID |
| FDR | in list | genes |  |  |
| 4.1e-06 | 22 | 1117 | Epithelium development | GO:0060429 |
| 1.0e-05 | 16 | 621 | Epithelial cell differentiation | GO:0030855 |
| 4.0e-05 | 37 | 3437 | Animal organ development | GO:0048513 |
| 4.9e-04 | 24 | 1855 | Tissue development | GO:0009888 |
| 5.3e-04 | 12 | 503 | Morphogenesis of an epithelium | GO:0002009 |
| 5.3e-04 | 12 | 500 | Gland development | GO:0048732 |
| 1.8e-03 | 7 | 167 | Mammary gland development | GO:0030879 |
| 2.1e-03 | 10 | 404 | Epithelial cell proliferation | GO:0050673 |
| 2.6e-03 | 39 | 4627 | System development | GO:0048731 |
| 2.6e-03 | 2 | 2 | Proximal/distal pattern formation involved in metanephric nephron development | GO:0072272 |
| GO-enrichment analysis of Stromal/Luminal/Alveolar cells of top 100 |  |  |  |  |

| genes found to be differentially expressed in samples of pregnant mice and animals with mammary gland involution |  |  |  |  |
| --- | --- | --- | --- | --- |
| Enrichment | Genes | Total | Functional Category | GO ID |
| FDR | in list | genes |  |  |
| 2.2e-04 | 36 | 3522 | Response to stress | GO:0006950 |
| 2.2e-04 | 12 | 448 | Response to oxidative stress | GO:0006979 |
| 2.2e-04 | 27 | 2100 | Cell death | GO:0008219 |
| 2.2e-04 | 28 | 2378 | Response to external stimulus | GO:0009605 |
| 2.2e-04 | 25 | 1949 | Programmed cell death | GO:0012501 |
| 4.5e-04 | 24 | 1909 | Apoptotic process | GO:0006915 |
| 4.6e-04 | 22 | 1654 | Positive regulation of molecular function | GO:0044093 |
| 5.4e-04 | 26 | 2247 | Catabolic process | GO:0009056 |
| 5.9e-04 | 24 | 1985 | Cellular catabolic process | GO:0044248 |
| 6.5e-04 | 22 | 1728 | Regulation of cell death | GO:0010941 |

The top 10 enriched categories are shown

Besides the examination of original expression profiles, we also made a comparison between the samples after the “style transfer” procedure: when samples of pregnant mice were transformed into a virgin or involution state, virgin – to pregnant or involution, involution – to pregnant or virgin. As an example, Fig. 5 shows MA-plots with comparison of Virgin versus Pregnant states of stromal cells (shown with blue dots), and Virgin versus artificial Virgin state created from Pregnant by style transfer (shown with orange). The overlay of these MA-plots provides a clear illustration that gene expression of original Virgin state is closer to that of artificially obtained Virgin than to original Pregnant samples.

**Fig. 5.**
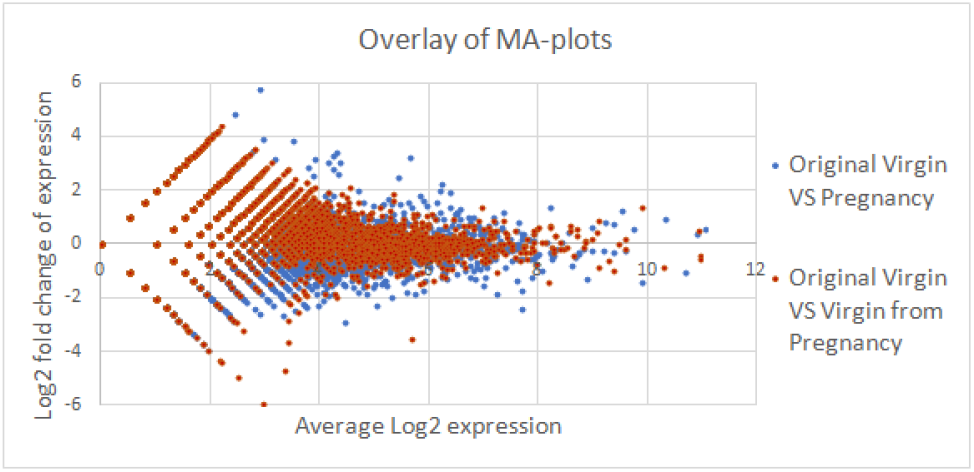
MA-plots comparing the gene expression in stromal cells from murine mammary glands in original and transformed samples. The comparison of original samples is shown with blue; the comparison between the original virgin state and the virgin state produced from pregnancy with style transfer is shown with orange color.

However, the similar GO- and KEGG-enrichment analysis of recovered gene expression and semisynthetic samples obtained with style transfer was less straightforward since there were numerous changes associated with basic GO categories. Thus, we decided to compare the variation of gene expression associated with relevant GO categories: mammary gland development (GO:0030879) and positive regulation of apoptotic process (GO:0043065). The highest variance in expression of genes involved in mammary gland development was observed in samples from pregnant mice (Fig. 6). The similar results were observed bo h in stromal and luminal cells (A) and also with using all the cells (B). The recovered expression was similar to original values, but the most interesting is that the style transfer from Virgin state to Involution and Pregnancy and from Involution to Virgin and Pregnancy resulted in biologically relevant changes in gene expression (the two lower panels of Fig. 6A and Fig. 6B).

**Fig. 6.**
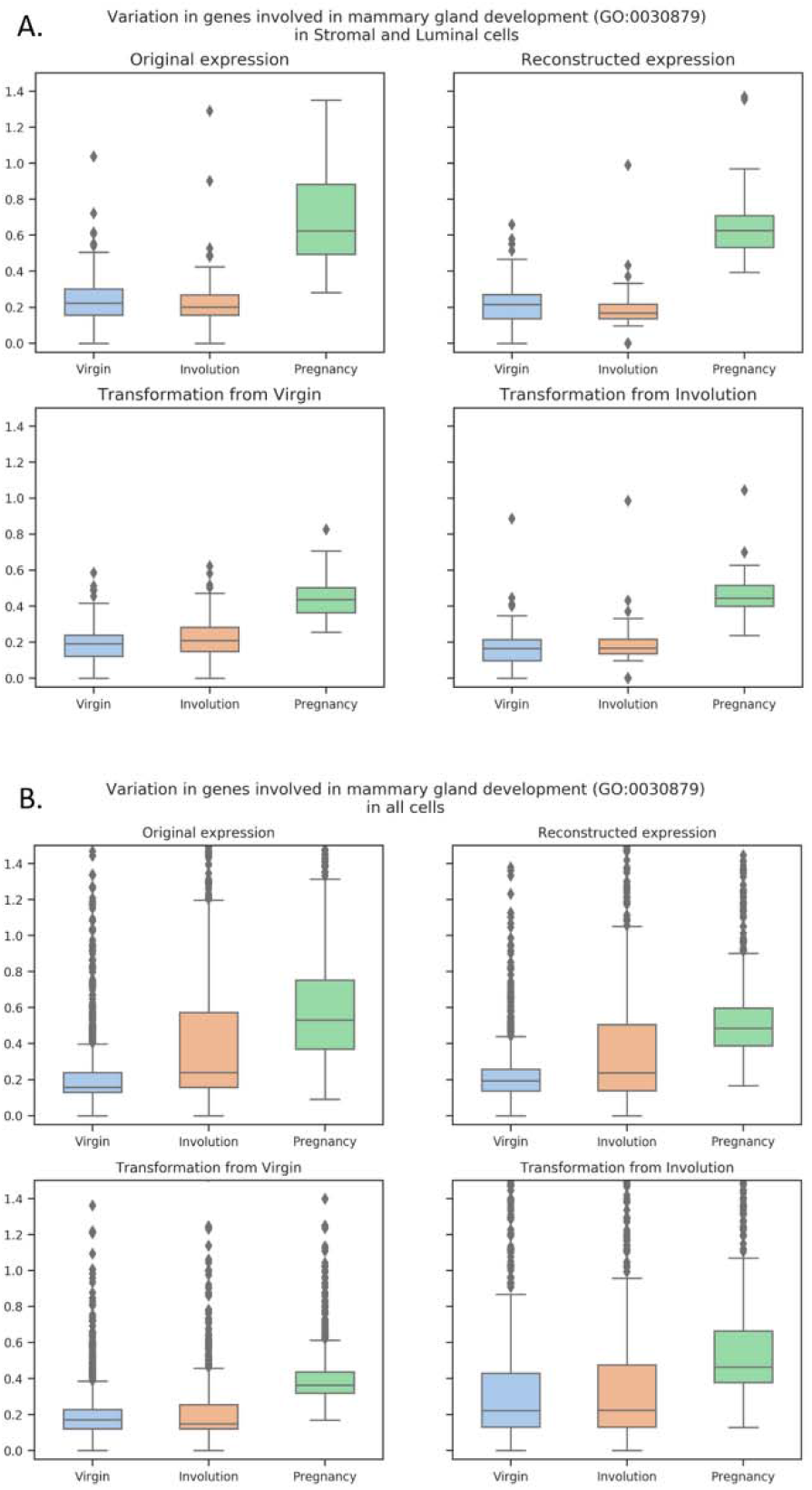
Variation in gene expression related to mammary gland development (GO:0030879) in Stromal and Luminal cells (A) and in all cells (B).

The similar analysis of genes involved in apoptosis regulation revealed two different pictures (Fig. 7). When only stromal, luminal and alveolar cells were considered the maximal variance was observed in samples from pregnant mice, and the second-high values were observed in samples from virgin mice (Fig. 7A, the top panels), however when all the cell types were considered the maximal variance was observed in samples from involuting mammary gland – as it was expected (Fig. 7B, the top panels). However, the results of style transfer (Fig. 7, the bottom panels) also demonstrate that the variance in apoptosis-related genes is higher in Involution state. The contradiction observed when only stromal, luminal and alveolar cells were considered might be due to the striking differences in proportions of various cell types. Thus, from here we can propose the additional advantage of style transfer procedure as it might be of help in studying gene expression changes resulting from certain biological or technical traits using the same initial data and treating the resulting samples as paired data.

**Fig. 7.**
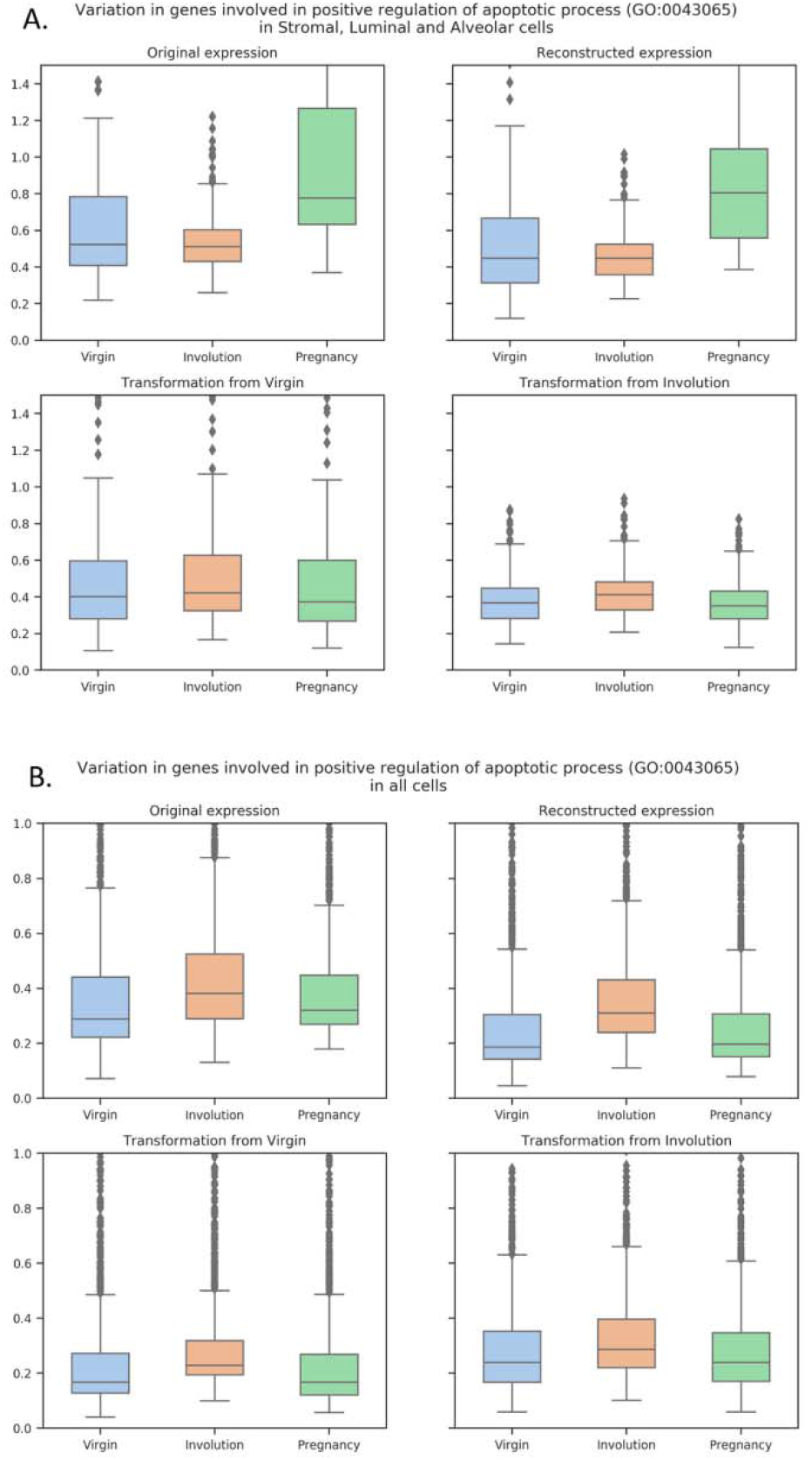
Variation in gene expression related to positive regulation of apoptotic process (GO:0043065) in Stromal, Luminal and Alveolar cells (A) and in all cells (B).

### 4.4 Biological examination of gene expression changes after encoding and decoding transformation using IFNβ-stimulated/unstimulated PBMCs scRNA-Seq data

The GSE96583 data was used to answer the question if style transfer can lead to biologically interpretable changes in predicted gene expression. The training subset was used to finetune the model and then all the analysis was performed on a validation subset. This dataset was also used to build the models with leave-one-group out approach: each time the whole group of particular cell type and condition was excluded from training. We have built four additional models: trained without stimulated CD4+ T-cells, unstimulated CD4+ T-cells, without either stimulated or unstimulated FCGR3A+ monocytes. The models were used for encoding/decoding the original samples with and without the style transfer. The obtained datasets were used for differential gene expression analysis. The top 100 differentially expressed genes were analyzed for GOenrichment. The results are partially presented on Fig. 8. In total, 66 pairwise comparisons were performed, the complete results can be found in supplementary table ST5. The “birds-eye” view of the table ST5 is shown on Fig. 8C. The yellow color indicates the sample pairs whose conditions were different either originally or after the style-transfer. The four colored columns of the table contain (1) the Jaccard index values characterizing the overlap between top 100 differentially expressed genes found in current samples with 87 genes found to be differentially expressed in original stimulated vs. control cells comparison (O(S) vs. O(C)) - chosen as etalon; (2) the numbers of significantly enriched GO categories; (3) the Jaccard index between the top 10 GO categories and top 10 GOs found in O(S) vs. O(C) comparison; and (4) the Jaccard index between the top 30 GO categories found in current DEGs and top 30 GOs in O(S) vs. O(C). The color changes from red to blue, which corresponds to a change in value from low to high. Generally, the most significantly enriched GO categories, observed in S vs. C comparisons, were associated with type I interferon response, as it was expected. Usually we saw no GO-enrichment in top differentially expressed genes if conditions of compared subsets were the same. Thus, the GO-enrichment analysis demonstrated that gene expression data after the style transfer contained relevant biological signals. The “expression changes” resulting from style transfer are further illustrated on Fig. 9, where the expression values of genes, associated with type I interferon signalling pathway (GO:0060337), are shown.

**Fig. 8.**
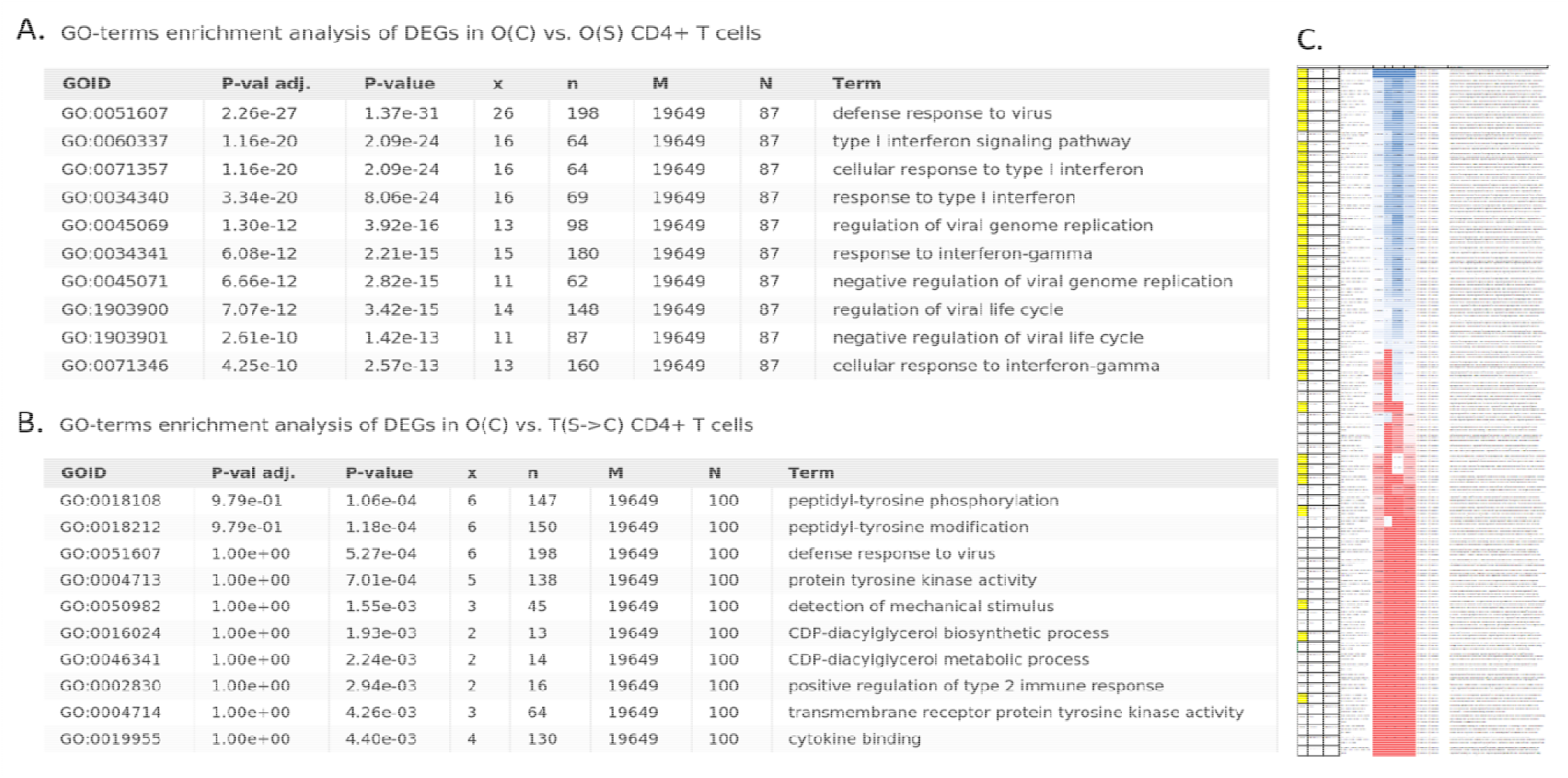
Results of GO-enrichment analysis of top 100 differentially expressed genes between different conditions. All the combinations of original and transfer conditions, models were tested – negative expression values were set to zero, then top 100 differentially expressed genes were selected using Welch’s test and analyzed for GO-terms enrichment. A. – GO-terms enrichment of genes differentially expressed in O(C) and O(S) CD4+ T-cells – from the original data; B. – GO-terms enrichment of genes differentially expressed in original unstimulated CD4+ T-cells – O(C) – and in T(S->C) – the interferon-treated CD4+ T-cells that were turned into unstimulated condition with our style-transfer procedure. C. – a “birds-eye” view of the table with results of all to all comparison in terms of differentially expressed genes overlap and GO-terms enrichment analysis. The yellow color indicates comparisons of samples whose conditions were different either originally or after the style-transfer. The four colored columns correspond to (1) the Jaccard index between top 100 differentially expressed genes found in current samples combination and in O(S) vs. O(C) (chosen as etalon); (2) a number of significantly enriched GO categories; (3) the Jaccard index between top 10 GO categories (sorted according to enrichment significance) against the top 10 GOs found in O(S) vs. O(C) DEG comparison; (4) the Jaccard index between top 30 GO categories and the top 30 GOs in O(S) vs. O(C) comparison. The color varies from red to blue corresponding to value changes from low to high. The original table can be found in supplementary table ST6.

**Fig. 9.**
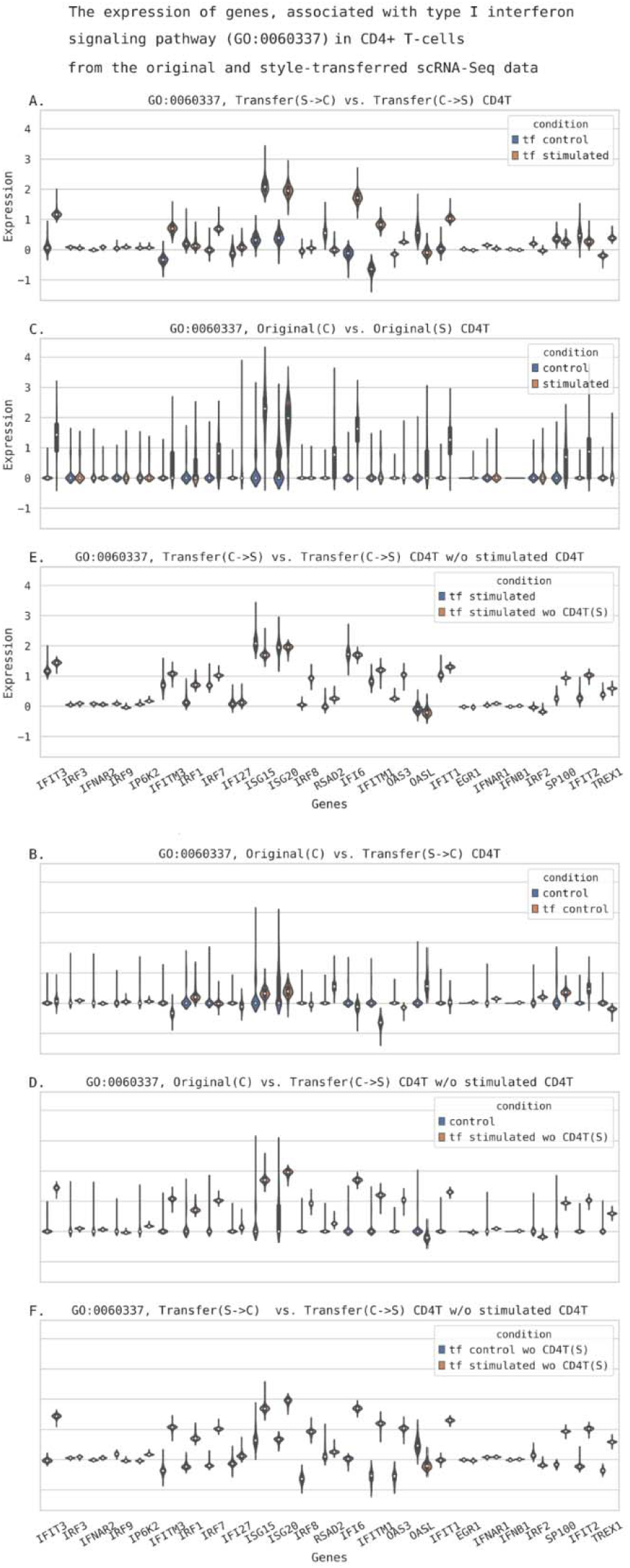
The original expression (decimal logarithm values) and the expression reconstructed from autoencoder after style-transfer in genes, associated with type I interferon signaling pathway, in CD4+ T cells. A. – the reconstructed gene expression in CD4+ T cells after transformation from interferon-stimulated state into unstimulated (control) state (S->C) and after the transformation from control to interferon-stimulated state (C->S); B. – the comparison of original gene expression in unstimulated CD4+ T cells – Original(C) – against the expression obtained with transformation from interferon-stimulated into control state; C. – the comparison of original gene expression in unstimulated – Original(C) – against stimulated CD4+ T cells – Original(S); D. – the original gene expression in unstimulated CD4+ T cells – Original(C) – against the style tra sfer C->S obtained with the model trained without the original stimulated CD4+ T cells gene expression data; E. – the transformations from control to stimulated state were done using a model pretrained on the whole training GSE96583 dataset (tf stimulated) or with a model trained on a subset of the training samples without stimulated CD4+ T cells (tf stimulated wo CD4T(S)); F. – the S->C and S->C transformations from a model trained without seeing the stimulated CD4+ T cells expression data. The validation subset of GSE96583 data was used.

The genes activity patterns in either reconstructed or transferred interferon-stimulated state resemble those observed in original data. Similar plots obtained for genes involved into two non-relevant biological processes - acetyl-CoA metabolic process (GO:0006084) and amyloid-beta metabolic process (GO:0050435) - demonstrated no alterations as compared to control. These figures (S3 and S4) can be found in Supplementary file SF2.

Pearson’s correlation coefficient of top 150 differentially expressed genes expression profiles between C->S transferred and O(S) samples was equal to 0.98, between S->C transferred and O(C) samples it was about 0.93 and for the sets with different conditions it was about 0.69-0.79. The higher conformity between the top differentially expressed genes in samples with corresponding states was illustrated with scatterplots on Figure S5 in Supplementary file SF2.

Thus, we can conclude that style-transfer approach can efficiently reconstruct gene expression changes associated with particular stimuli. With control CD4+ T cells transformation into interferon-stimulated condition we observed relevant changes in gene expression and GO-enrichment.

The semisynthetic samples became similar to original interferon-stimulated cells. Switching the state of stimulated cells into untreated condition resulted in decreased expression of genes involved into interferon response pathways. It is also of interest that the models built with leave-one-group-out approach were found to be reasonably accurate. Similar findings were also observed with other cell subsets (data not shown).

## 5 Discussion

Construction of information-rich, low-dimensional representations of gene expression profiles remains a challenging task. Availability of such representations is a gateway to successful data harmonization, domain adaptation and deeper understanding of interconnections between expression of various genes. The proposed framework allows to investigate gene expression profile shifts when some specific, pre-defined categorical factor of variation changes. The framework performs the dimensionality reduction of gene expression data in such a way that hidden variables are disentangled into two separate domains where one subgroup is fully interpretable and accounted for chosen, pre-defined factors of variation and another, larger group of hidden variables is designed to contain no information about these factors. So that, we can controllably change the factor(s) and see the impact on the gene expression level. This leads us to several possibilities: either to harmonize the data by using the batch and style codes (either biological traits, e.g. some treatment condition or cell type, or technical, e.g. sequencer model) as factors of style and to perform the downstream analysis on the latent variables (which also dramatically reduces the dimensionality) instead of raw expression, or to transfer selected samples to a particular style trying to predict the effect of desired treatment/phenotype combination in silico. Ability to obta n gene expression values of (semi)synthetic samples makes it possible to analyze the predicted changes using the well-accepted and interpretable bioinformatic approaches, for example to check which genes became the most differentially expressed when particular technical or biological traits were changed with style transfer, which GO-categories or molecular pathways were predicted to be affected the most etc. The proposed approach can help to solve different problems associated with real transcriptomic data, e.g., to reduce the variance associated with batch effects, to check the data for outliers, to reduce the data dimensionality retaining relevant biological information.

Our future efforts on the framework will be mostly conducted towar s increasing the fidelity of generated samples and further evaluation of our approach on different datasets and comparing its performance with existing frameworks. Also, future research will include the induction of sparsity in both encoder and decoder weights to figure out its effect on framework performance in terms of variation sources disentanglement and applicability of generated samples to downstream bioinformatic pipelines. Moreover, the sparse weights promise some insights on what genes affect each other expression and affected by choice of the style.

## Acknowledgements

Authors would like to thank the Institute of Computational Technologies SB RAS for providing computational resources needed for this publication.

## Conflict of Interest

none declared.

## Notes

https://github.com/NRshka/stvae-source

https://doi.org/10.6084/m9.figshare.9925115

### Competing Interest Statement

The authors have declared no competing interest.

### Summary of Updates

We have substantially updated the manuscript, performed the benchmarking with several similar tools and performed additional in silico studies

## References

Amodio, M. et al. (2019). Exploring single-cell data with deep multitasking neural networks. Nat Methods, 16, 1139–1145.

Bult,C.J. et al. (2019) Mouse Genome Database (MGD). Nucleic Acids Res., 47, D801–D806.

Collado-Torres,L. et al. (2017) recount workflow: Accessing over 70,000 human RNA-seq samples with Bioconductor. F1000Research, 6, 1558.

Eraslan,G. et al. (2019) Single-cell RNA-seq denoising using a deep count autoencoder. Nat. Commun., 10, 390.

Gatys,L.A. et al. (2015) A Neural Algorithm of Artistic Style. arXiv, 1508.06576.

Ge,S.X. and Jung,D. (2018) ShinyGO: a graphical enrichment tool for animals and plants. bioRxiv, 315150.

Ghahramani,A. et al. (2018). Generative adversarial networks simulate gene expression and predict perturbations in single cells. bioRxiv, 10.1101/262501.

Gold,M.P. et al. (2018) Shallow Sparsely-Connected Autoencoders for Gene Set Projection. In, Biocomputing 2019. WORLD SCIENTIFIC, pp. 374–385.

Grønbech,C.H. et al. (2018) scVAE: Variational auto-encoders for single-cell gene expression data. bioRxiv, 318295.

Han,X. et al. (2018) Mapping the Mouse Cell Atlas by Microwell-Seq. Cell, 172, 1091–1107.e17.

Higgins,I. et al. (2017) beta-VAE: Learning Basic Visual Concepts with a Constrained Variational Framework. In, ICLR.

Hoffman,J. et al. (2017) CyCADA: Cycle-Consistent Adversarial Domain Adaptation.

Johansen,N. and Quon,G. (2019) scAlign: a tool for alignment, integration, and rare cell identification from scRNA-seq data. Genome Biol., 20, 166.

Lachmann,A. et al. (2018) Massive mining of publicly available RNA-seq data from human and mouse. Nat. Commun., 9, 1366.

Liu,L. et al. (2019) On the Variance of the Adaptive Learning Rate and Beyond.

Lopez,R., et al. (2018) Deep generative modeling for single-cell transcriptomics. Nat Methods. 15, 1053–1058.

Lotfollahi,M., et al. (2019) scGen predicts single-cell perturbation responses. Nat Methods 16, 715–721 (2019).

Lotfollahi,M., et.al. (2019). Conditional out-of-sample generation for unpaired data using trVAE. ArXiv,1910.01791.

Mescheder,L.M. et al. (2018) Which Training Methods for GANs do actually Converge? arXiv, 1801.04406.

Misra, D. (2019). Mish: A Self Regularized Non-Monotonic Neural Activation Function. arXiv, 1908.08681.

Patacchiola,M. et al. (2019). Y-Autoencoders: disentangling latent representations via sequential-encoding. arXiv, 1907.10949.

Romanov,A. et al. (2018) Adversarial Decomposition of Text Represe tation. arXiv, 1808.09042.

Saliba,A.-E. et al. (2014) Single-cell RNA-seq: advances and future challenges. Nucleic Acids Res., 42, 8845–8860.

Shekhar,K. et al. (2016) Comprehensive classification of retinal bipolar neurons by single-cell transcriptomics. Cell, 166, 1308–1323.

Sohn,K. et al. (2015) Learning structured output representation using deep conditional generative models. In Proceedings of the 28th International Conference on Neural Information Processing Systems - Volume 2 (NIPS’15). MIT Press, Cambridge, MA, USA, 3483–3491.

Stark,R. et al. (2019) RNA sequencing: the teenage years. Nat. Rev. Genet. 20, 631–656.

Targonski,C., et al. (2019). Cellular State Transformations using Generative Adversarial Networks. arXiv, 1907.00118.

Wang,D. & Gu,J. (2018). VASC: Dimension Reduction and Visualization of Single-cell RNA-seq Data by Deep Variational Autoencoder. Genomics, Proteomics & Bioinformatics. 16. 320–331.

Wang,X. et al. (2018). Three-dimensional intact-tissue sequencing of single-cell transcriptional states. Science, 27, 361(6400), pii: eaat5691..

Way,G.P. and Greene,C.S. (2018) Extracting a biologically relevant latent space from cancer transcriptomes with variational autoencoders. Biocomput. 2018, 80–91.

Xu,C. et al. (2019) Harmonization and Annotation of Single-cell Transcriptomics data with Deep Generative Models. bioRxiv, 532895.

Zheng,G.X.Y. et al. (2017) Massively parallel digital transcriptional profiling of single cells. Nat. Commun. 8, 14049.

Zhu,J.-Y. et al. (2017) Unpaired Image-to-Image Translation using Cycle-Consistent Adversarial Networks. arXiv, 1703.10593.

Ziemann,M. et al. (2019) Digital expression explorer 2: a repository of uniformly processed RNA sequencing data. Gigascience, 8, pii: giz022.

